# Architectural asymmetry enables DNA transport through the *Helicobacter pylori cag* type IV secretion system

**DOI:** 10.1101/2023.07.25.550604

**Authors:** Mackenzie E. Ryan, Prashant P. Damke, Caitlynn Bryant, Michael J. Sheedlo, Carrie L. Shaffer

## Abstract

Structural asymmetry within secretion system architecture is fundamentally important for apparatus diversification and biological function. However, the mechanism by which symmetry mismatch contributes to nanomachine assembly and interkingdom effector translocation are undefined. Here, we show that architectural asymmetry orchestrates dynamic substrate selection and enables trans-kingdom DNA conjugation through the *Helicobacter pylori cag* type IV secretion system (*cag* T4SS). Structural analyses of asymmetric units within the *cag* T4SS periplasmic ring complex (PRC) revealed intermolecular π-π stacking interactions that coordinate DNA binding and license trans-kingdom conjugation without disrupting the translocation of protein and peptidoglycan effector molecules. Additionally, we identified a novel proximal translocation channel gating mechanism that regulates cargo loading and governs substrate transport across the outer membrane. We thus propose a model whereby the organization and geometry of architectural symmetry mismatch exposes π−π interfaces within the PRC to facilitate DNA transit through the *cag* T4SS translocation channel.

## INTRODUCTION

Architectural symmetry mismatch is a remarkable evolutionary innovation exhibited by diverse bacterial nanomachines including the dynamic type IV secretion system (T4SS) superfamily^1–3^. Recent advances in the structural definition of paradigmatic T4SS machineries uncovered unexpected asymmetric features at the junction between the inner membrane complex (IMC) and outer membrane complex (OMC), or between concentric rings comprising the OMC in expanded T4SS architectures^1–4^. In addition to T4SS machineries, symmetry mismatch has been observed in other bacterial secretion systems including the T2SS^5^, T3SS^6, 7^, and T6SS^8^, suggesting an important, yet unresolved, functional significance. Although the structural relationships between T4SS apparatus components and cognate effector substrates transiting the secretion channel are currently unresolved, sub-structure symmetry mismatch presumably provides conformational mobility to facilitate the unidirectional ratcheting of selected cargo between adjacent architectural layers. Accordingly, asymmetry between structural sub-complexes may impart the incredible functional plasticity and apparatus dynamism required to orchestrate diverse substrate selection and enable directional effector translocation^1, 3, 9^.

The versatile T4SS nanomachine is employed by Gram-negative and Gram-positive organisms to deliver DNA, proteins, or other substrates to target prokaryotic or eukaryotic cells. In Gram-negative organisms, ‘minimized’ T4SS machineries encompass roughly 12 conserved components, denoted VirB1 through VirB11 and VirD4, following a standardized nomenclature derived from the prototypical *Agrobacterium tumefaciens vir* T4SS^10–12^. In these streamlined configurations, the apparatus IMC subassembly forms part of the translocation channel that spans the periplasm and couples to the OMC through a stalk-like structure^9, 13–15^. In some architectures, elongated VirB10 domains act as connectors bridging the OMC and IMC^9, 11, 15^. Multiple ATPases, including the VirD4 coupling protein that delivers nucleoprotein complexes to the secretion channel^11, 16–19^, power cargo translocation across the bacterial envelope.

Trans-kingdom DNA transfer achieved by minimized T4SS machineries is an evolutionary remnant of generalized interbacterial conjugation mechanisms^10, 20, 21^. Thus far, relatively few ‘dual function’ systems that deliver nucleoprotein complexes across kingdom boundaries have been identified. The prototypical dual function *vir* T4SS harbored by *A. tumefaciens* translocates tumor-inducing DNA (T-DNA) and associated effector proteins into recipient plant cells in a contact-dependent manner^10, 20, 22–25^. Seminal work using a transfer DNA immunoprecipitation (TrIP) assay^24^ to trace the DNA translocation pathway through the *vir* T4SS secretion channel provided a mechanistic understanding of how apparatus components coordinate nucleoprotein delivery across the bacterial outer membrane. In addition to *A. tumefaciens*, recent studies demonstrate that dual purpose and expanded machineries in *Rhizobium etli*^26^, *Bartonella henselae*^27–30^, *Coxiella burnetii*^29^, *Legionella pneumophila*^29, 31, 32^, and *Helicobacter pylori*^33^ have the fundamental capacity to translocate DNA into eukaryotic cells via T4SS-dependent mechanisms.

An emerging paradigm for understanding how a single molecular device transports diverse immunostimulatory cargo into eukaryotic cells, the cancer-associated *H. pylori cag* T4SS translocates the bacterial oncoprotein CagA and an expanded repertoire of DNA, LPS metabolite, and polysaccharide substrates into gastric epithelial cells. In contrast to the *A. tumefaciens vir* T4SS that delivers plasmid-borne single-stranded DNA (ssDNA) into plant cells, the *H. pylori cag* T4SS delivers double-stranded chromosomal fragments (dsDNA) into epithelial cells via unresolved mechanisms^21, 33, 34^. Multiple lines of evidence demonstrate that *cag* T4SS-dependent DNA translocation is susceptible to exogenous DNA-targeting nucleases and anti-DNA monoclonal antibodies^33, 34^, suggesting a two-step translocation mechanism that does not appear to require a conjugative pilus conduit. *H. pylori* trans-kingdom DNA conjugation elicits both anti- and pro-inflammatory responses via TLR9 stimulation^21, 33, 35^ and activation of additional nucleic acid reconnaissance systems^34^. Consequently, *H. pylori* evolved mechanisms to counterbalance nucleic acid surveillance and suppress STING signaling within the gastric mucosa^35, 36^. Thus, *cag* T4SS-dependent DNA translocation provides additional mechanisms by which *H. pylori* manipulates the immune response to enable persistent gastric niche colonization^33, 35^.

Similar to other ‘expanded’ systems, the *cag* T4SS apparatus is comprised of an IMC harboring a trio of ATPases connected to the OMC via a structurally unresolved stalk-like region^17, 37^. Within the OMC, the outer membrane-associated cap incorporates the structural components CagX, CagY, CagT, CagM, and Cag3, and includes a periplasmic ring complex (PRC) comprised of portions of CagX and CagY^1, 3, 17, 37, 38^. Analysis of the *cag* T4SS cryo-EM structure revealed striking symmetry mismatch whereby several CagX subunits in the PRC (14-fold symmetry) do not display an obvious connection to corresponding subunits within the OMC (17-fold symmetry)^3^. Asymmetry between OMC subassemblies is observed in other ‘expanded’ systems including the *Legionella pneumophila dot/icm* T4SS^39^ and conjugative F-plasmid machinery^40, 41^. Although architectural asymmetry is a common feature among macromolecular complexes, the biological significance of symmetry mismatch between T4SS subassemblies remains unknown.

Here, we employed a structure-function approach to delineate mechanisms underscoring *cag* T4SS-dependent DNA translocation. We report the identification of a novel *cag* T4SS gating mechanism that regulates the passage of DNA and protein effectors across the outer membrane and licenses the import of exogenous molecules into the periplasm. We provide evidence that in addition to direct interactions with components of the *cag* T4SS IMC and PRC, the VirB2-like pilin CagC polymerizes to form an ‘endopilus’ gate or plug domain that controls effector transport through the secretion channel. We propose a model in which CagC polymers dynamically assemble into an endopilus-like structure corresponding to the stalk region that bridges the IMC and periplasmic complexes in the *cag* T4SS machinery. Mechanistically, we demonstrate that the VirB9 ortholog CagX facilitates secondary substrate selection within the PRC via dsDNA-π−stacked interactions at sites exhibiting architectural asymmetry. Whereas disruption of intermolecular CagXπ-π stacking resulted in a significant reduction of DNA binding *in vitro* and diminished trans-kingdom conjugation *in vivo*, non-nucleic acid cargo was effectively delivered into gastric epithelial cells when PRC π-stacking interactions were genetically ablated. Collectively, these studies highlight architectural innovations underpinning substrate selectivity and provide a mechanistic framework for understanding how the remarkable *cag* T4SS nanomachine coordinates diverse effector molecule delivery into gastric epithelial cells.

## RESULTS

### CagC gates effector translocation through the *cag* T4SS

The *cag* T4SS harbors multiple *vir* T4SS homologs including CagX, CagY, CagT, CagV, CagE, Cagα, CagL, CagC, and Cag5 (VirB9, VirB10, VirB7, VirB8, VirB4, VirB11, VirB5, VirB2, and VirD4, respectively)^1, 3, 17, 18, 22, 38, 42–44^. Prior work identified ‘uncoupling’ mutations within the *A. tumefaciens vir* T4SS that selectively disarm T-pilus biogenesis without perturbing trans-kingdom conjugation^12, 45, 46^. Similarly, various *H. pylori cag* T4SS-dependent phenotypes, including CagA translocation and IL-8 stimulation, occur via uncoupled mechanisms that require incongruent subsets of *cag* T4SS components^18, 34, 47, 48^, raising the hypothesis that molecularly dissimilar *cag* T4SS substrates are secreted into host cells via distinct mechanisms. To test this hypothesis, we first analyzed the contribution of *vir* T4SS homologs in TLR9 stimulation, an outcome that requires a functional *cag* T4SS^18, 33, 34, 49^. Consistent with previous studies analyzing *cag* T4SS-dependent phenotypes^18, 35, 43, 49^, TLR9 stimulation occurred in a CagA-independent manner^33, 34^ and required recognized *vir* T4SS homologs, with the exception of the VirD4-like coupling protein Cag5^18^, suggesting that DNA trafficking through the secretion channel occurs via non-canonical trans-kingdom conjugation (**Fig. 1a**). To investigate the mechanism by which disparate *cag* T4SS substrates are selected for transport to target host cells, we next analyzed potential structural features that could facilitate DNA substrate selection for export through the *cag* T4SS apparatus. Previous work identified a single point mutation within the *A. tumefaciens vir* T4SS OMC that confers secretion channel gating defects resulting in the uncontrolled release of VirE2 effectors to the bacterial cell surface^50^. We reasoned that the *cag* T4SS may harbor a similar gating mechanism based on the presence of multiple *vir* T4SS homologs localized throughout the apparatus architecture^1, 3, 18, 38^. To test whether individual *cag* T4SS components are associated with apparatus gating, we employed non-denaturing colony blot assays developed to interrogate translocation channel integrity in *A. tumefaciens*^50^. In contrast to WT *H. pylori*, which does not release detectable levels of *cag* T4SS effector molecules into the extracellular milieu, isogenic mutants deficient in *cagC* accumulated surface-exposed CagA and transfer DNA in a host cell contact-independent manner (**Fig. 1b**). Genetic complementation of the *cagC* isogenic mutant at a heterologous chromosomal locus abrogated effector leakage (**Fig. 1b**) and restored *cag* T4SS-dependent TLR9 activation (**Fig. 1c**) and IL-8 stimulation phenotypes (**Fig. 1d**), suggesting that CagC regulates channel gating or substrate passage across the outer membrane. In support of the hypothesis that CagC forms a specialized plug or gating mechanism, inactivation of other reported structural (CagX, CagY, CagT, CagV, CagL) or energetic (CagE, Cagα, Cag5) components did not affect *cag* T4SS translocation channel integrity (**Fig. 1b**).

**Figure 1.**
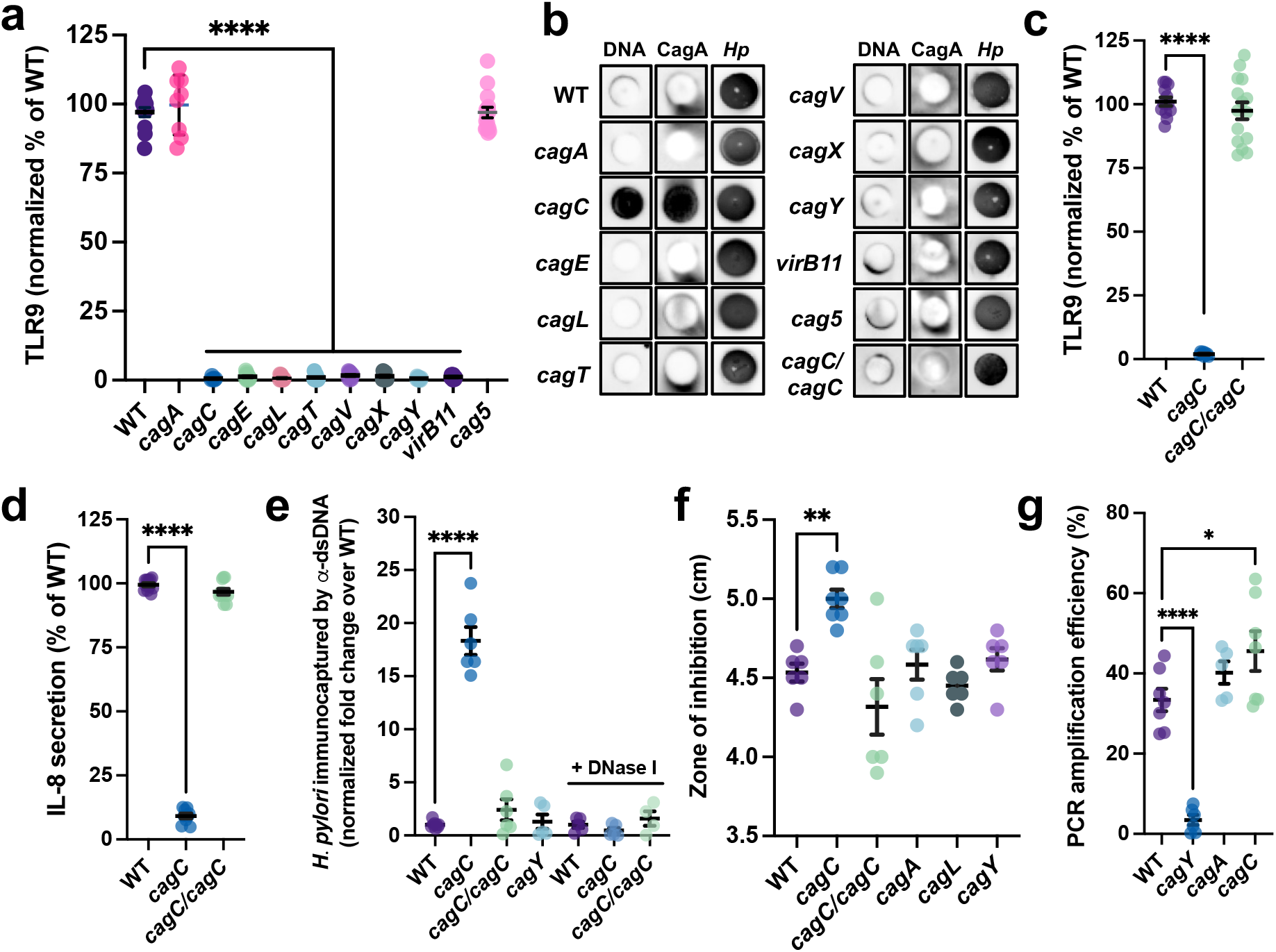
CagC gates the *cag* T4SS translocation channel. (**a**) Levels of *cag* T4SS-dependent TLR9 activation induced by the indicated *H. pylori* 26695 isogenic mutant strain. Data are expressed as the normalized fold change over mock infected cells as a percent of TLR9 stimulation achieved by WT. (**b**) Non-denaturing colony blots probing surface-exposed dsDNA, CagA, and *H. pylori* (*Hp*) antigens. (**c**) TLR9 stimulation and (**d**) IL-8 secretion requires *cagC*. (**e**) Levels of viable *H. pylori* immunocaptured by anti-dsDNA monoclonal antibodies. Data are expressed as the normalized fold change in recovered colony forming units over levels obtained for WT. (**f**) Erythromycin susceptibility of the indicted strain. Data depict the zone of inhibition for the indicated strain. (**g**) Transfer DNA immunopurification assays demonstrating the presence of *H. pylori* chromosomal DNA fragments within the *cag* T4SS apparatus. Graph depicts the amplification efficiency of a 795 bp fragment in transfer DNA assay preparations purified from the indicated strain. Amplification efficiency of immunopurified samples is expressed as a percent of the input DNA amplification from at least four biological replicate experiments. In **a** and **c**-**g**, significance was determined by one-way ANOVA with Dunnett’s post-hoc correction for multiple comparisons to experimental controls; *, *p*<0.05, **, *p*<0.01, ****, *p*<0.0001.

To quantitatively monitor the uncontrolled release of effector DNA into the extracellular milieu, we developed a live cell assay to enumerate *H. pylori* that accumulate surface-exposed transfer DNA in the absence of host cell contact. For these studies, WT *H. pylori* or the indicated isogenic mutant strain was incubated with anti-double-stranded DNA (anti-dsDNA) monoclonal antibodies or an equivalent volume of a corresponding IgG isotype control, and immunocaptured bacteria were isolated by Protein G-conjugated magnetic bead separation. Compared to WT and the *cagY* strain, which does not assemble the *cag* T4SS core complex^1, 3, 38^ and thus does not exhibit effector leakage (**Fig. 1b**), significantly more *cagC* bacteria were immunocaptured by anti-dsDNA antibodies (**Fig. 1e**). Consistent with a role in effector gating, genetic complementation of the *cagC* mutant restored immunocaptured bacteria to WT levels (**Fig. 1e**). Likewise, incubating bacterial cultures with DNase I prior to anti-dsDNA immunocapture markedly reduced levels of isolated *H. pylori cagC* (**Fig. 1e**) demonstrating that aberrantly localized dsDNA is tethered to the bacterial cell surface.

We next sought to determine whether *cag* T4SS pore gating defects afford unrestricted access to the periplasm. We reasoned that loss of either secretion channel integrity or pore gating would license the influx of bulky molecules, such as erythromycin, that are typically impermeable to the Gram-negative outer membrane^50–52^. To test whether potential *cag* T4SS pore gating mechanisms control periplasm access, we employed erythromycin sensitivity assays to detect changes in outer membrane permeability^51^. Compared to WT and isogenic mutants that do not exhibit effector molecule leakage, the *cagC* strain displayed significantly increased pore permeability (**Fig. 1f**) indicating that in the absence of host cell contact, CagC is important for stabilizing the secretion channel in a tightly closed conformation. Erythromycin susceptibility patterns of the genetically complemented *cagC* mutant phenocopied the parental WT strain (**Fig. 1f**), further demonstrating that CagC controls pore gating and access to the periplasm via leaky secretion channels.

Previous work determined that effector DNA is pre-loaded into the *cag* T4SS apparatus prior to initiating host cell contact to generate a ‘ready-to-fire’ nanomachine^34^. To test whether defective gating modulates DNA loading, we used a modified transfer DNA immunoprecipitation assay developed in *A. tumefaciens*^24^ to monitor levels of DNA confined within the *cag* T4SS core complex lumen^34^. For these studies, we isolated *cag* T4SS assemblies from chemically cross-linked *H. pylori* via immunopurification targeting CagY (a constituent of the *cag* T4SS outer membrane-associated complex and periplasmic ring structure^1, 3, 38^), and monitored DNA loading by PCR analysis of co-purifying chromosomal fragments that remained shielded from DNase degradation within the secretion channel. In agreement with a previous report^34^, DNA fragments were readily amplified from cross-linked CagY preparations obtained from WT and *cagA*-deficient *H. pylori*, but not from mock preparations generated from the *cagY* isogenic strain (**Fig. 1g**). Consistent with a role in regulating DNA passage through the *cag* T4SS OMC, significantly more DNA co-purified with CagY complexes in strains lacking *cagC* (**Fig. 1g**), demonstrating that CagC is not required for effector DNA loading into the distal secretion channel. Collectively, these studies demonstrate that loss of CagC confers *cag* T4SS channel leakiness that enables the unrestricted export of effectors and the import of exogenous small molecules into the periplasm.

### CagC is a specialized T4SS pilin ortholog

We next performed protein sequence analysis and structural modeling to further investigate the role of CagC as a novel gating mechanism within the *cag* T4SS apparatus. In accordance with a previous study analyzing pilin-like motifs in bacterial cell surface-exposed polypeptides^53^, CagC exhibited primary sequence similarity to canonical T4SS pilins including *A. tumefaciens* VirB2 and *E. coli* TraA (**Supplemental Fig. 1a**). AlphaFold2^54^ modeling of the mature CagC monomer lacking the leader peptide^54–56^ revealed a predicted helical structure that closely resembles the experimentally resolved structures of conjugative pilins VirB2^57, 58^ and TraA^59^ (**Fig. 2a** and **Supplementary Fig. 1b,c**). In contrast to a previous report suggesting that conjugative pilins are cyclic in nature^60^, superimposition of predicted CagC structures with monomeric VirB2 and TraA revealed a slightly U-shaped protomer corresponding to structural conformations in which the free pilin termini were oriented towards the exterior of assembled canonical T-pilus or F-pilus filaments^57–59^ (**Fig. 2a**).

**Figure 2.**
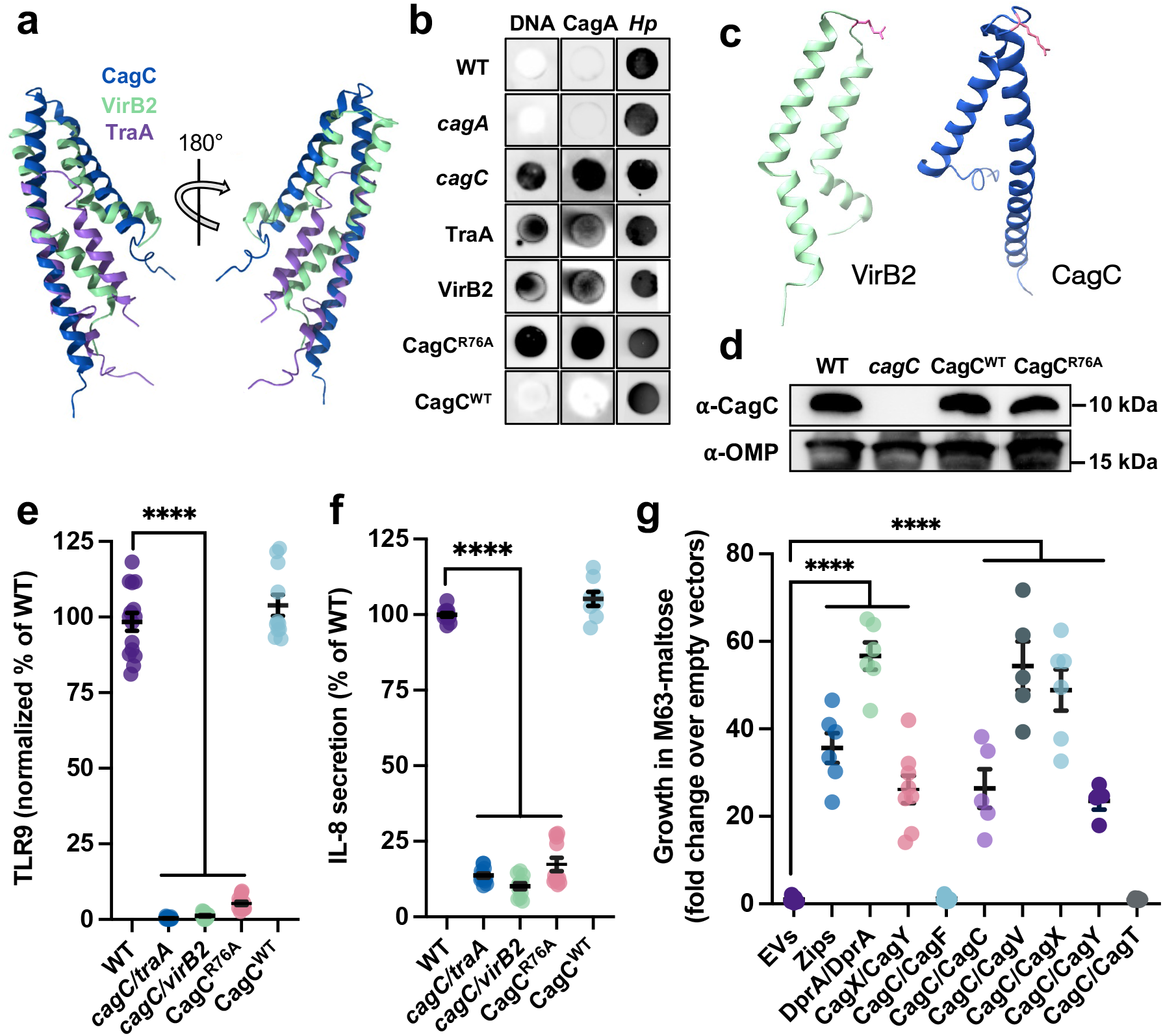
CagC is a VirB2-like T4SS pilin ortholog. (**a**) Superimposed model of CagC, VirB2 (T-pilus, PDB: 8CUE), and TraA (F-pilus, PDB: 5LEG). Structures of mature CagC were predicted using AlphaFold2, and the highest-ranked structure was superimposed with canonical pilin structures using the Matchmaker function in ChimeraX. (**b**) Colony blots demonstrating dsDNA and CagA cell surface localization in *H. pylori cagC* mutants complemented by either *traA, virB2,* or the CagC^R76A^ variant. (**c**) Structures of VirB2 (PDB: 8CUE) and CagC (AlphaFold2 modeling) highlighting Arg 91 (VirB2) and Arg 76 (CagC) residues in the pilin luminal loop region (pink coloring). (**d**) Western blot analysis of *H. pylori cagC* mutants genetically complemented by CagC^WT^ or the CagC^R76A^ variant. OMP, outer membrane protein. (**e**) Levels of TLR9 stimulation and (**f**) IL-8 secretion induced by the indicated strain. (**g**) Bacterial two-hybrid analysis of CagC protein-protein interactions. Data depict the fold change in the OD_600_ of *E. coli* BTH101 cultures transformed with the indicated plasmid pairs and propagated in M63-maltose. Positive protein-protein interactions are indicated by growth in M63-maltose and data are expressed as the fold change in growth over BTH101 transformed with empty vectors (EVs). In **a**, **e** and **f**, significance was determined by one-way ANOVA with Dunnett’s post-hoc correction for multiple comparisons to experimental controls; ****, *p*<0.0001.

Based on the predicted structural conservation, we hypothesized that orthologous pilins could rescue *cag* T4SS defects exhibited by the *cagC* isogenic mutant. To these this hypothesis, we expressed either *A. tumefaciens virB2* or *E. coli traA* in the *cagC* isogenic mutant and analyzed *cag* T4SS-dependent outcomes elicited by each chimera. Despite predicted structural similarities, genetic complementation of *cagC* with either *virB2* or *traA* did not restore *cag* T4SS pore gating (**Fig. 2b**), TLR9 activation (**Fig. 2e**), or IL-8 stimulation (**Fig. 2f**) defects, suggesting that CagC fulfills a unique role within the *cag* T4SS. One possibility is that CagC forms an ‘endopilus’ within the secretion channel lumen corresponding to the stalk-like density observed in the *in situ cag* T4SS architecture^17, 37^. Recent studies analyzing VirB2 protomers identified arginine residues (Arg 91) that protrude into the T-pilus lumen and form extensive electrostatic interactions required for pilus assembly and structural stabilization^57, 58^ (**Fig. 2c**). Congruent with a predicted function as a specialized pilin, CagC harbors a single arginine residue in the mature protomer (Arg 76) corresponding to VirB2 Arg 91 (**Fig. 2c**). We reasoned that similar to T-pilus biogenesis, polymerization of a putative ‘endopilus’ could be selectively disrupted by modifying positively charged residues within the predicted CagC luminal loop (**Fig. 2a,c**). To investigate whether Arg 76 is required for CagC function, we generated a CagC variant containing a R76A single point mutation expressed from a heterologous chromosomal locus in the *cagC* mutant. Western blot analysis of *H. pylori* whole cell lysates reveled high levels of CagC^R76A^ expression (**Fig. 2d**), demonstrating that in contrast to VirB2 Arg 91^57, 58^, CagC Arg 76 does not contribute to protomer stability. Consistent with a role as a putative ‘endopilus’ or gating mechanism, CagC^R76A^ exhibited uncontrolled DNA and effector protein release to the bacterial cell surface (**Fig. 2b**). Accordingly, compared to the parental WT strain and *cagC* mutants genetically complemented with CagC^WT^, CagC^R76A^ variants were defective in TLR9 activation (**Fig. 2e**) and IL-8 stimulation (**Fig. 2f**), indicating that positively charged residues within the luminal loop are important for CagC protomer function or multimerization into higher order structures. Taken together, these data suggest that the specialized pilin CagC assembles into a novel plug or gate-like apparatus to control effector passage through the secretion system pore.

### Periplasmic *cag* T4SS assemblies bind transfer DNA

We next sought to understand the mechanism by which CagC regulates effector transit through the translocation channel. In the mature form, CagC pilin subunits are approximately 9.2 kDa (**Fig. 2d**) and attempts to generate a functional epitope-tagged fusion protein for immunopurification studies were unsuccessful. Therefore, we employed a bacterial two-hybrid (BTH) screening approach designed to characterize interactions among both soluble and membrane-associated components^61–63^ to localize CagC complexes within the *cag* T4SS apparatus. Analysis of direct protein-protein interactions revealed that in comparison to negative (empty vector) and positive (Zip-Zip^62^ and *H. pylori* DprA-DprA^64^) controls, CagC strongly interacted with components predicted to comprise the T4SS apparatus inner membrane complex (IMC) including the bitopic VirB8 homolog CagV (**Fig. 2g**). BTH assays also revealed strong interactions among the CagX N-terminal domain (residues 41–310) and corresponding CagY fragments (residues 1469–1603, designated CagY_CT_) that assemble into the periplasmic ring complex (PRC)^1, 3, 38^, demonstrating the ability of our screening approach to detect direct protein-protein interactions occurring in the periplasm. In addition to CagC-CagV, we detected CagC-CagC protomer interactions (**Fig. 2g**), suggesting that CagC can self-polymerize into higher order structures or subcomplexes. Whereas interactions among CagC and components localized to either the cytoplasm (the chaperone-like molecule CagF^65^) or the periphery of the *cag* T4SS outer membrane complex (the VirB7 homolog CagT^1, 3, 38^) were not detected, CagC directly interacted with both CagX and CagY_CT_ (**Fig. 2g**). These observations led to the hypothesis that CagC subassemblies bridge the IMC and PRC.

One possibility is that a putative CagC ‘endopilus’ assembly corresponding to the *cag* T4SS stalk-like region visualized *in situ*^17, 37^ may dynamically assemble and retract to regulate effector DNA gating within the PRC via a piston-like mechanism. To test the hypothesis that CagC interacts with DNA, we purified recombinant CagC and performed electromobility shift assays (EMSA) to analyze DNA binding capacities. In agreement with a previous study suggesting that dsDNA is delivered to host cells in a *cag* T4SS-dependent manner^34^, CagC weakly bound 58-mer effector dsDNA sequences^34^, but not corresponding 58-mer ssDNA targets (**Fig. 3a**). Based on the observation that CagC interacts with both CagX and fragments of CagY that comprise the PRC (**Fig. 2g and Fig. 3b,c**), we questioned whether other *cag* T4SS subcomplexes bind DNA to facilitate cargo transfer. Compared to maltose-binding protein (MBP) or recombinant MBP-CagT controls, MBP-CagX robustly bound target *H. pylori* DNA sequences^34^ in a concentration-dependent manner (**Fig. 3d,e** and **Supplemental Fig. 2a**). In contrast to CagX, TraK (*E. coli* F plasmid) and VirB9 (*A. tumefaciens vir* T4SS) homologs harbored by conjugative DNA transfer systems did not interact with DNA (**Supplemental Fig. 2b,c**), suggesting that DNA binding within the PRC is a unique structural innovation in the *cag* T4SS apparatus. Consistent with the observation that CagC displays preferential interactions with dsDNA substrates (**Fig. 3a**), recombinant CagX exhibited a stronger binding affinity for dsDNA compared to corresponding ssDNA sequences (**Fig. 3e**). In support of this result, DNA competition assays demonstrated CagX preferential binding to dsDNA (**Fig. 3f** and **Supplemental Fig. 2d-f**). Together, these studies identify CagX as a novel periplasmic DNA-binding T4SS component.

**Figure 3.**
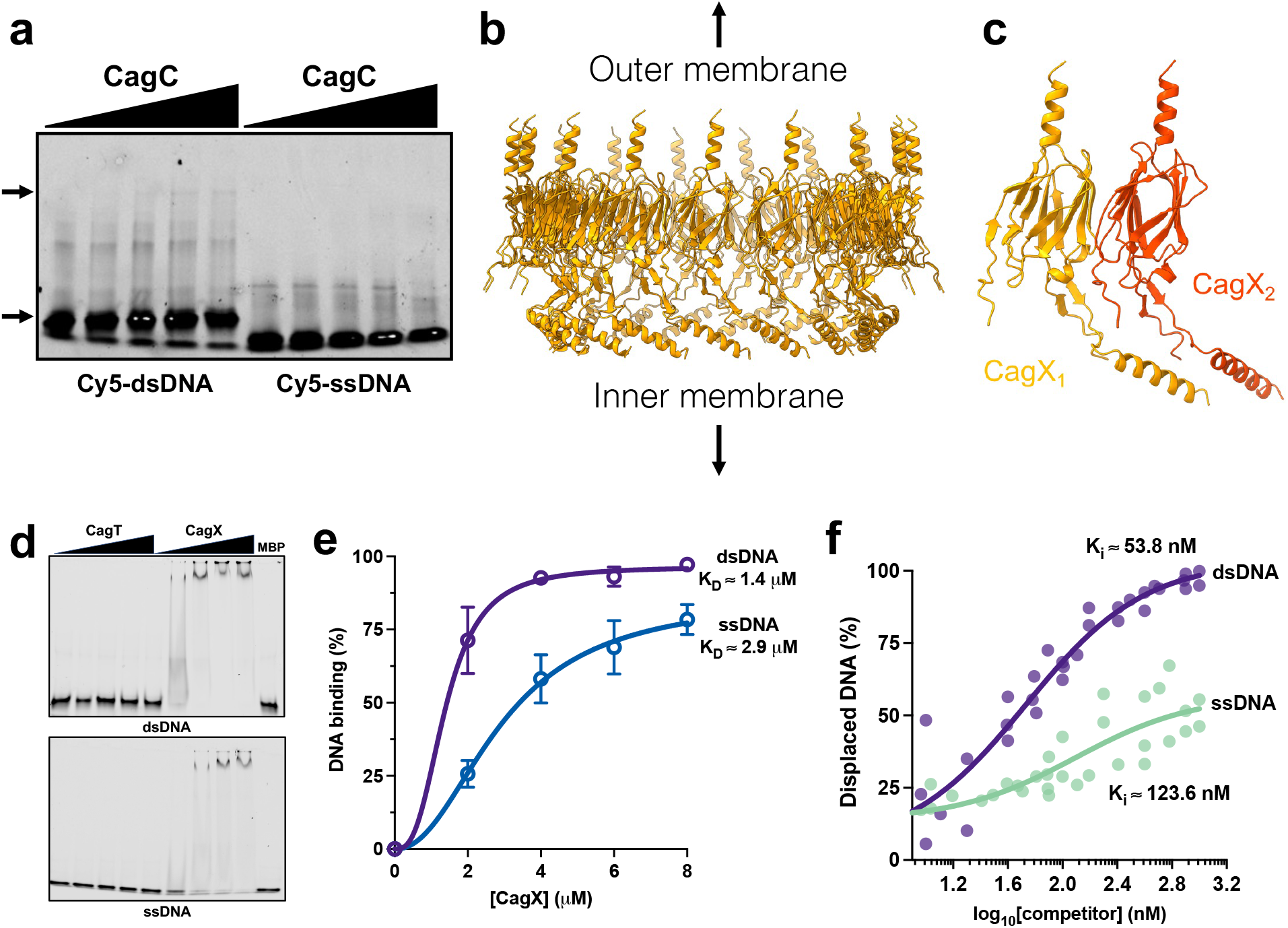
Asymmetric units within the *cag* T4SS PRC bind dsDNA. (**a**) EMSA analysis of recombinant MBP-CagC binding to 58-mer dsDNA (left) or ssDNA (right). Arrows indicate recombinant CagC-DNA complexes. (**b**) The position of CagX subunits within the PRC (PDB: 6X6J). (**c**) Structure of adjacent CagX subunits (CagX_NT_, residues 41-310) within the assembled PRC. (**d**) EMSA analysis of recombinant CagX binding to dsDNA (upper panel) and ssDNA (lower panel). MBP-CagX, but not MBP or MBP-CagT, binds target dsDNA and ssDNA. (**e**) Quantitation of MBP-CagX binding to the indicated DNA substrate. Values indicate the experimentally approximated K_D_ for CagX-DNA binding to 58-mer targets. (**f**) Competition assays demonstrating preferential CagX binding to dsDNA. Data points represent the % displaced DNA obtained in independent biological replicate experiments. Solid line represents the nonlinear regression estimating the inhibitory constants (K_i_) for dsDNA and ssDNA. In **e**, dissociation constants (K_D_) were estimated using the specific binding with hill slope nonlinear regression model and in **f**, inhibitory constants (K_i_) were estimated using the one site fit K_i_ model (heterologous ligand binding affinities using experimentally derived K_D_ in **e**) in GraphPad Prism 9.5.

### Symmetry mismatch enables DNA loading into the secretion channel

To delineate the molecular determinants of DNA binding within the *cag* T4SS PRC, we employed a structure-guided analysis to identify potential DNA binding sites in CagX-CagY assemblies. Close inspection of the CagX structure^1, 3^ (**Fig. 3c**) revealed a basic patch formed at the interface of adjacent CagX subunits within the PRC (**Fig. 4a**) that represented a potential DNA binding region. We hypothesized that overlapping π (pi) orbitals among a series of arginine (Arg 36, 38, 278, and 295) and tryptophan (Trp 58) residues in the basic patch would produce electrostatic interfaces that enable DNA interactions within the secretion channel (**Fig. 4b**). Notably, the diameter and geometry of the proposed DNA binding site supports this assignment (**Fig. 4c**). To test the hypothesis that electrostatic CagX interfaces facilitate DNA loading into the secretion channel, we generated CagX variants by introducing point mutations in adjacent arginine residues within the basic patch region to disrupt intermolecular π-stacking interactions (**Fig. 4b**). In comparison to CagX^WT^, CagX variants in which Arg 36 and Arg 38 were replaced with alanine (CagX^R36A,R38A^) exhibited similar dsDNA binding affinities *in vitro*, whereas CagX variants harboring the Arg 278 single point mutation (CagX^R278A^) exhibited significantly decreased DNA binding capacities that were further reduced when consecutive Arg residues were disrupted in tandem (CagX^R36A,R38A,R278A^) (**Fig. 4d** and **Supplemental Fig. 3a,b**). Congruent with our hypothesis, *H. pylori cagX* mutants expressing CagX^R36A,R38A^, CagX^R278A^, or CagX^R36A,R38A,R278A^ translocated significantly less DNA into host cells, resulting in markedly reduced levels of TLR9 activation compared to the parental WT strain and *cagX* mutants genetically complemented by CagX^WT^ (**Fig. 4e**). Analysis of additional *cag* T4SS-dependent phenotypes produced by the CagX^R36A,R38A,^, CagX^R278A^, and CagX^R36A,R38A,R278A^ variant strains revealed that despite the observed DNA translocation defects, levels of IL-8 secretion were comparable to the WT and CagX^WT^ complemented strains (**Fig. 4f**), suggesting that DNA delivery is mechanistically uncoupled from the translocation of other *cag* T4SS substrates. In contrast, disrupting Arg 295 via single point mutation or in tandem with Arg 36, Arg 38, or Arg 278 destabilized CagX, resulting in significantly decreased DNA binding *in vitro* (**Supplementary Fig. 3b,c**) and markedly reduced *cag* T4SS activity (**Supplementary Fig. 3d-f).** We thus speculate that Arg 295 constitutes a critical CagY-binding interface in the PRC. Collectively, these studies indicate that intermolecular CagX π-stacking interactions generate a DNA-binding pocket required for periplasmic substrate selection and trans-kingdom conjugation.

**Figure 4.**
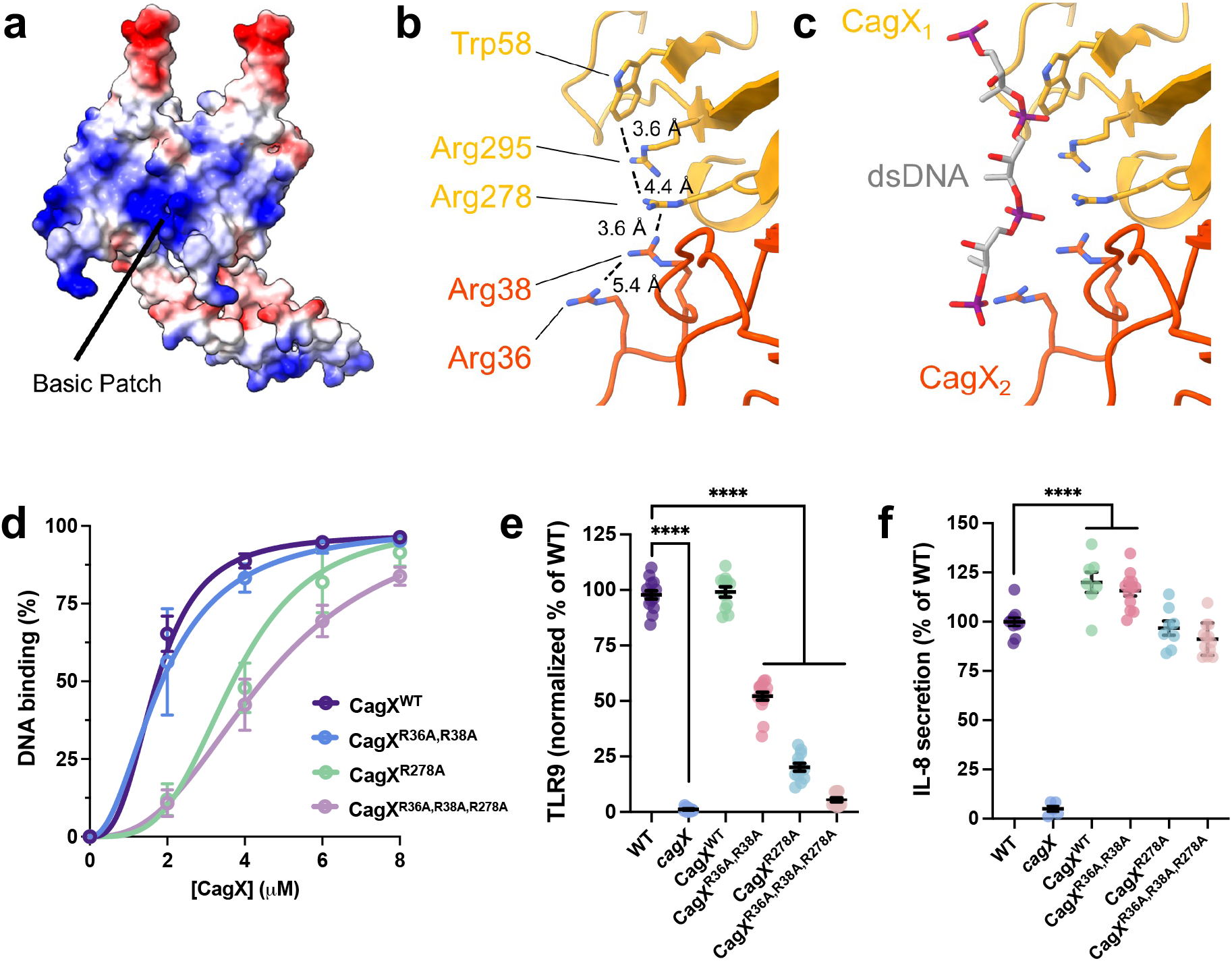
Symmetry mismatch coordinates DNA substrate selection for trans-kingdom conjugation. (**a**) Electrostatic surface potential of two adjacent CagX residues demonstrating the formation of a basic patch at the intersubunit junction. Regions of acidic and basic potential are shown in red and blue, respectively. (**b**) Overlapping π−π orbital interactions among CagX arginine and tryptophan residues in adjacent subunits. (**c**) Proposed model of dsDNA binding to CagX subassemblies. (**d**) Quantitation of CagX variant binding to dsDNA. Experimentally approximated K_D_ for CagX^WT^ (1.7 µM), Cag^XR36A,R38A^ (1.9 µM), CagX^R278A^ (3.7 µM), and CagX^R36A,R38A,R278A^ (4.6 µM) were calculated for 58-mer target binding. (**e**) Levels of TLR9 activation and (**f**) IL-8 secretion stimulated by *H. pylori* harboring the indicated CagX variant. In **d**, dissociation constants (K_D_) were estimated using the specific binding with hill slope nonlinear regression model in GraphPad Prism 9.5. In **e** and **f**, significance was determined by one-way ANOVA with Dunnett’s post-hoc correction for multiple comparisons to experimental controls; ****, *p*<0.0001.

Analysis of *cag* T4SS architecture resolved by single particle reconstruction indicates that portions of CagX and CagY assemble into structures localized to both the OMC and the PRC (**Fig. 5**), producing a striking symmetry mismatch^1, 3^. In the current study, we identified a putative DNA binding site on CagX that is centered around resides Arg 36, 38, 278, 295 and Trp 58. The high-resolution cryo-EM structure indicates that this site is occupied by the periplasmic portion of CagY and thus, DNA binding may be excluded in the assembled PRC^3^. However, we note that the cryo-EM structure was determined through the imposition of symmetry and argue that this map omits asymmetric features. To that end, a low-resolution reconstruction of the PRC detected 14 copies of CagX in the OMC and 17 copies of CagX in the PRC (**Fig. 5a**) with 14 tube-like helical densities bridging the OMC (CagX C-terminus) and the PRC (CagX N-terminus)^3^ (**Fig. 5b,c**). This observation suggests that the additional copies of CagX in the PRC provide DNA interaction sites to facilitate nucleic acid trafficking through the core complex (**Fig. 5d**). We thus propose a model whereby asymmetric regions of the PRC assemble to form a DNA binding site for substrate selection and loading into the *cag* T4SS distal secretion channel (**Fig. 5d**).

**Figure 5.**
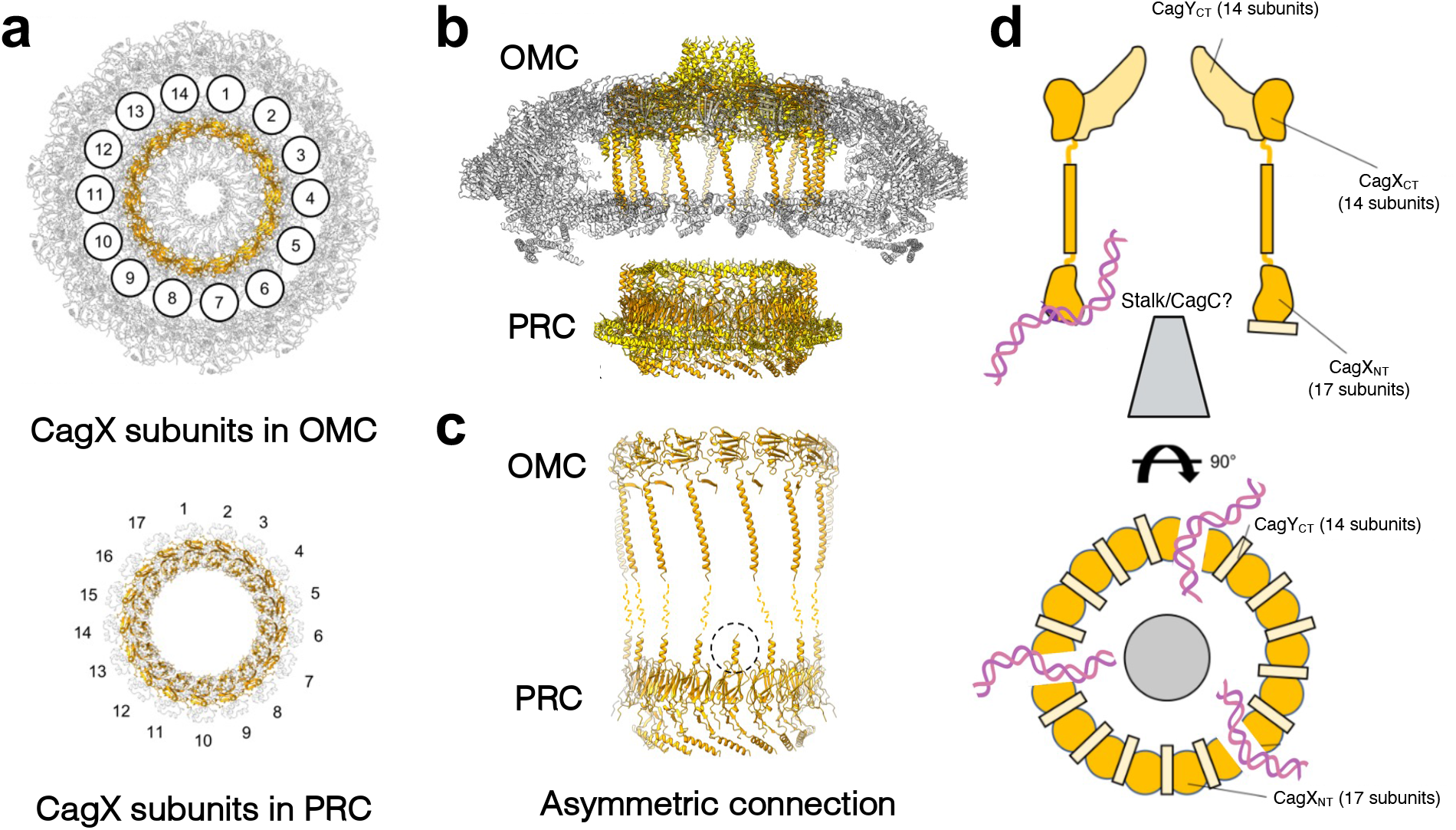
Proposed model of DNA substrate selection within the *cag* T4SS PRC. (**a**) Symmetry mismatch between CagX subunits in the OMC (upper panel, 14-fold symmetry) and the PRC (lower panel, 17-fold symmetry) single particle cryo-EM reconstruction. (**b**) Positions of CagY (yellow) and CagX (orange) within the *cag* T4SS OMC and the PRC. (**c**) Illustration of one asymmetric CagX connection between the OMC and PRC (dashed circle). Three copies of CagX in the PRC do not show an obvious connection to corresponding CagX subunits in the OMC. (**d**) Proposed model of dsDNA loading into the *cag* T4SS core complex via electrostatic interactions and π−π stacking within asymmetric PRC units. Illustration depicts the architectural juxtaposition of CagX N-terminal domains (CagX_NT_), CagX C-terminal domains (CagX_CT_), CagY N-terminal domains (CagY_NT_), and CagY C-terminal domains (CagY_CT_) in the OMC and PRC with dsDNA loading into the secretion channel via sub-complex symmetry mismatch.

## DISCUSSION

Trans-kingdom DNA conjugation is an ancient mechanism that profoundly impacts host cell biology. The identification of DNA substrate selection mechanisms within the *cag* T4SS PRC highlights novel evolutionary adaptations that enable expanded machinery function. As previously suggested for the *A. tumefaciens vir* T4SS^66^, the VirB9 ortholog CagX exhibits N-terminal domains that contribute to protein stability and establish inter-subunit contacts required for DNA substrate selection and trafficking through the translocation channel. Notably, our study uncovered multiple ‘uncoupling’ mutations that specifically disrupt DNA translocation without perturbing delivery of other substrates into host cells, highlighting remarkable *cag* T4SS apparatus flexibility. The capacity of CagX to recognize DNA and mediate selective substrate transfer through the *cag* T4SS OMC provides intriguing evidence suggesting that in addition to prototypical coupling proteins, periplasmic architectural features recruit effector cargo to the secretion channel. We speculate that CagX architectural asymmetry thus represents a secondary substrate selection checkpoint that regulates cargo transit through the periplasm and across the distal OMC translocation channel. We hypothesize that symmetry mismatch in other dual function T4SS machineries, and potentially other diverse secretion system architectures, similarly orchestrates substrate selection to ensure spatiotemporal and unidirectional translocation into target cells. Collectively, these observations expand the growing collection of disengaged systems exhibiting selective substrate translocation^34, 47^ through evolutionarily divergent T4SS nanomachines in which scaffold biogenesis is uncoupled from trans-kingdom DNA conjugation^50, 66, 67^.

Our studies to define the architectural determinants of *cag* T4SS substrate selection revealed a novel CagC-dependent gating mechanism that regulates cargo transfer through the proximal translocation channel into the PRC (**Fig. 1,2**). Consistent with a proposed role as an endopilus- like structure, previous work demonstrates that CagC is a membrane-associated VirB2 ortholog that is localized to the bacterial cell surface via a process that requires *cag* T4SS apparatus biogenesis^53^. Additional studies report the assembly of filamentous structures at the *H. pylori*- gastric epithelial cell interface that were proposed to constitute an extracellular *cag* T4SS- associated appendage analogous to interbacterial conjugative pili or the *A. tumefaciens* T pilus^48, 53, 68–70^. However, the molecular composition of these structures has not been elucidated and multiple reports have identified several inconsistencies that argue against the presence of a canonical *cag* T4SS conjugative pilus. For example, *H. pylori* isogenic mutants deficient in the OMC component CagY, which forms the central translocation channel and is required for core complex assembly^1, 3, 4, 38^, produce extracellular filaments at the same level as WT bacteria^68, 69^. Thus, it is difficult to envision how a *cag* T4SS-associated pilus would nucleate and polymerize in the absence of (i) a translocation channel required for pilin transport to the bacterial cell surface and (ii) the OMC platform on which the conjugative pilus assembles^71^. While a putative *cag* T4SS- associated pilus may nucleate in the periplasm or at the inner membrane complex via mechanisms similar to T3SS needle biogenesis^72^, evidence strongly supports a model whereby prototypical T4SS conjugative pilus structures polymerize at the outer membrane^71^.

Furthermore, extracellular *cag* T4SS-associated filaments have not been observed in frozen hydrated *H. pylori*-gastric cell co-cultures imaged via cryo-electron tomography (cryo-ET) under near native conditions^17, 37^. While a conjugative pilus was not directly observed in the current study, our results provide compelling evidence for the assignment of CagC as a VirB2 pilin structural ortholog that forms a gating mechanism within the proximal secretion channel. Similar to the conjugative F pilus and the *vir* T4SS T pilus, we propose that CagC protomers assemble into a pentameric endopilus-like structure that lines the secretion channel. Assembled CagC pentamers may serve as a scaffold for biogenesis of heteromeric complexes that incorporate the proposed VirB5-like minor pilin CagL^70, 73, 74^ to form the cone-shaped stalk density that bridges the IMC and PRC^17, 37^. Stalk-like structures with similar sizes and geometries have been identified in cryo-ET atomic models of other T4SS machineries including the R388 conjugative system^9^ and the *L. pneumophila dot/icm* T4SS^75–77^. Alternatively, CagC may form the rod structure observed spanning the periplasm and originating from inner membrane platforms within the *cag* T4SS architecture *in situ* ^37^. Finally, in conjunction with the observation that *H. pylori* deficient in *cagC* produce filamentous structures at the microbe-epithelial cell interface^69^, the discovery of outer membrane-derived nanotubes produced by *H. pylori* in both the presence and absence of host cell contact^37, 78^ support the hypothesis that the intercellular structures visualized in scanning electron microscopy studies are bacterial cell surface appendages unrelated to the *cag* T4SS.

*H. pylori* trans-kingdom conjugation occurs via incompletely defined mechanisms that require evolutionarily divergent VirD2-like relaxases and the DNA translocase/partitioning protein FtsK for nucleic acid cargo coupling to the translocation channel^34^. Our work points towards a novel two-step substrate translocation mechanism that involves secondary DNA substrate selection within asymmetric CagX subunits in the PRC (**Fig. 5**). In this model, membrane tethered FtsK may form a pore in the inner membrane to pump effector DNA into the periplasmic space for substrate selection by the *cag* T4SS PRC. Alternatively, FtsK may act as a modified coupling protein that displaces cognate Cag5 from the *cag* T4SS cytoplasmic apparatus to deliver DNA directly into the proximal secretion channel. In this scenario, the stalk-like structure may dynamically retract and extend to ensure efficient DNA loading into the distal translocation channel. Secondary DNA substrate selection within the PRC chamber would therefore safeguard unidirectional translocation into host cells. This observation is strikingly similar to the previously suggested models of DNA trafficking through the *A. tumefaciens vir* T4SS whereby VirB9 C-terminal domains presumably interact with cellular VirB2 pilin subunits to stabilize the secretion channel^24, 66^. Likewise, the identification of VirB10-dependent gating mechanisms within the interior of the *vir* T4SS OMC chamber^50^ and single point mutations in CagC that license the uncontrolled release of *cag* T4SS substrates to the bacterial cell surface (**Fig. 1,2**) suggest that translocation channel gating is a fundamental mechanism within diverse dual function T4SS architectures. In the context of previously identified *vir* T4SS ‘uncoupling’ mutations that enable DNA translocation in the absence of extracellular pilus biogenesis, and evidence demonstrating that *cag* T4SS-dependent DNA transfer is susceptible to exogenous nucleases and neutralizing DNA antibodies^33, 34^, we speculate that CagC forms an internal endopilus that directs DNA through the distal secretion channel. We propose that while CagC is not required for DNA loading into the proximal translocation channel (**Fig. 1g**), putative endopilus-like structures may trigger conformational changes that stabilize the OMC pore. Subsequently, CagC polymers may line the translocation channel lumen to form a tight mating junction spanning the cellular interface. As CagC is required for the delivery of all identified *cag* T4SS substrates across the host cell membrane^53, 69^ (**Fig. 1**), a putative CagC endopilus may function as a target cell sensor that locks the OMC pore into a translocation-competent state or as a signal transducer that propagates external cues to stimulate substrate switching. Coupled to secondary substrate selection mechanisms within the *cag* T4SS PRC, these studies broaden our understanding of how nanomachine architectural asymmetry orchestrates the release of diverse molecular cargo across the bacterial outer membrane. Future work to decipher complex structural relationships within the assembled secretion channel and periplasmic apparatus will undoubtedly provide mechanistic details underscoring interkingdom cargo transport through evolutionarily diverse machineries.

## MATERIALS AND METHODS

### Bacterial strains and culture conditions

*Helicobacter pylori* 26695 and isogenic derivatives were grown on trypticase soy agar plates supplemented with 5% sheep blood (BD) at 37°C with 5% CO_2_. Overnight cultures of *H. pylori* were grown in Brucella broth supplemented with 5% fetal bovine serum (FBS) at 37°C in 5% CO_2_. *E. coli* strain DH5α (New England Biolabs) was used for plasmid propagation and was grown on lysogeny broth (LB) agar or liquid media supplemented with the appropriate antibiotics for plasmid maintenance. Bacterial two-hybrid (BACTH) strain BTH101 (a gift from Scot Ouellette, University of Nebraska Medical Center) was maintained on LB, and BACTH screening was performed using LB plates supplemented with 0.5 mM isopropyl β-D-1-thiogalactopyranoside (IPTG), 100 μg/mL ampicillin, 50 μg/mL kanamycin, and 20 μg/mL 5-bromo-4-chloro-3-indoyl-β-D-galactopyranoside (X-Gal)^62^. BACTH-dependent growth selection was performed in M63 minimal media [(NH_4_)_2_SO_4_ (2 g/L), KH_2_PO_4_ (13.6 g/L), Thiamine B1 (1 mg/L), 1 mM MgSO_4_, FeSO_4_.7H_2_O (5 mg/L), pH 7] supplemented with 0.4% maltose, 0.5 mM IPTG, 50 µg/mL ampicillin, and 15 µg/mL kanamycin. All BACTH experiments were executed at 30°C under constant aeration.

### *H. pylori* mutagenesis

Isogenic mutants^18, 38, 48, 69^ were complemented in *cis* at the *ureA* locus using plasmids derived from pAD1^48^ that were designed to express either the native gene or a protein variant as previously described^48, 49^. Point mutants were generated using Q5 Site Directed Mutagenesis (New England Biolabs). Plasmid sequences were confirmed by PCR and whole plasmid sequencing, and constructs were used to transform isogenic mutant strains. Colonies resistant to kanamycin (12.5 μg/ml) or chloramphenicol (10 μg/mL) were selected and complementation at the *ureA* locus was confirmed by PCR and by anti-CagX immunoblotting, when appropriate.

### Human cell culture

HEK293-hTLR9 (Invivogen hkb-htlr9) and the corresponding parental HEK293 null1 (InvivoGen hkb-null1) cell lines were grown in DMEM media supplemented with 10% heat-inactivated FBS and 1X GlutaMAX (Life Technologies) in the presence of 5% CO_2_ at 37°C. AGS human gastric epithelial cells (ATCC CRL-1739) were grown in RPMI media supplemented with 10% FBS, 2 mM L-glutamine, and 10 mM HEPES in the presence of 5% CO_2_ at 37°C.

### TLR9 stimulation assay

TLR9 activation assays were carried out as previous described^35, 49^. Briefly, HEK293 cells expressing human TLR9 (HEK293-hTLR9) or the parental HEK293 null1 cells were co-cultured with WT *H. pylori* 26695 or *cag* mutant strains at a MOI of 100. Supernatants were collected at 24 hours post-infection, and TLR9 activation was quantified by measuring secreted embryonic alkaline phosphatase (SEAP) by QuantiBlue^TM^ reagent (Invivogen) using a microplate reader (BioTek Synergy HI) to record the absorbance at 650 nm. TLR9 activation was normalized to levels of SEAP produced by the corresponding infected null1 cells, and data were expressed as a percent of the normalized fold change over WT-challenged cells.

### Colony immunoblotting

To assess bacterial cell surface localization of *cag* T4SS effectors^50^, *H. pylori* strains were grown on blood agar plates for 24 h before harvesting and normalization to OD_600_ of 1. Bacteria were pelleted by centrifugation and resuspended in 50 μL Brucella broth to reach an OD_600_ of 20. Concentrated suspensions were spotted onto fresh blood agar plates (5 μL) and incubated at 37°C, 5% CO_2_. After 24 h, a nitrocellulose membrane was placed on top of the colonies for 5 min at room temperature. The membrane was washed twice in TBST (tris-buffered saline, 0.05% Tween 20) and blocked in 5% milk in TBST. Surface exposure of bacterial effectors was assessed by immunoblotting using anti-CagA (α-CagA, Santa Cruz Biotechnology) or anti-dsDNA (α- dsDNA, EMD Millipore). Immunoblotting with anti-*H. pylori* antibody (a gift from Dr. Tim Cover) was performed as a positive control.

### Live cell bacterial immunocapture

*H. pylori* was grown for 24 h on blood agar plates prior to collection and normalization to an OD_600_ of 0.5 in Brucella broth supplemented with 6% FBS. Normalized bacteria were incubated with 250 μg/mL anti-dsDNA monoclonal antibody (α-dsDNA, EMD Millipore) or an equivalent amount of IgG isotype control antibody (Thermo) for 1 h in the presence of 5% CO_2_ at 37°C. After incubation, an aliquot was obtained for total colony forming unit (CFU) enumeration by serial dilution plating. The remaining sample was incubated with Protein G Dynabeads (Invitrogen) for 10 minutes at RT with constant rotation. Beads were isolated by magnetic separation and washed twice with 5 bead volumes of sterile PBS. Beads were then resuspended in 100 μL sterile PBS followed by serial dilution and plating for CFU enumeration. For DNase treated samples, *H. pylori* pellets were resuspended in PBS and normalized to an OD_600_ of 1 prior to incubation with 5 μL DNase buffer and 1 unit Turbo DNase I (Life Technologies). After 15 min incubation, an equal volume of Brucella broth supplemented with 6% FBS was added to each well to reach an OD_600_ of 0.5, and bacterial immunocapture was performed. Levels of *H. pylori* displaying dsDNA on the bacterial cell surface were calculated as a percentage of total bacteria in each sample and normalized to levels of bacteria isolated in IgG isotype control purifications.

### Erythromycin susceptibility assays

*H. pylori* strains cultured for 24 h on blood agar plates were collected, normalized to an OD_600_ of 1, and an equivalent volume of normalized cultures were inoculated onto fresh blood agar by sterile bead plating. Erythromycin-impregnated discs (15 μg) were placed onto dried plates, and the diameter of the zone of inhibition was determined after 24 h incubation at 37°C and 5% CO_2_. Susceptibility assays were performed a minimum of three times for each *H. pylori* strain.

### ‘Transfer DNA’ immunoprecipitation

Transfer DNA assays were performed as previously described^24, 34^. Briefly, *H. pylori* protein-DNA complexes were cross-linked by the addition of 500 μL 1% paraformaldehyde followed by incubation for 10 min at room temperature and quenching with 1 ml 250 mM glycine. Cross-linked cells were lysed by sonication in lysis buffer (5 mM EDTA, 1% NP-40 in PBS) supplemented with 2X cOmplete™ inhibitor (Roche). Cell lysates were treated with 2 units of Turbo DNase (Life Technologies) and 10 mM MgCl_2_ for 30 min at room temperature. Solubilized membranes were separated by centrifugation at 14,000 rpm for 30 minutes at 4°C. CagY complexes were purified using polyclonal anti-CagY antisera (a gift from Dr. Tim Cover) conjugated to Protein G Dynabeads (Invitrogen) isolated by magnetic separation. Purified complexes were washed twice in PBS supplemented with 10 mM MgCl_2_ and 1 unit Turbo DNase, followed by two washes in high salt buffer (lysis buffer containing 400 mM NaCl), and a final wash in PBS. Following extensive washing, protein-DNA complexes were eluted and de-cross-linked in 100 µl 1% SDS in 0.1 M NaHCO_3_ at 65°C overnight. Proteins were digested using Proteinase K (10 µg) at 65°C for 30 minutes, and DNA was precipitated by 100% ethanol in 0.3 M sodium acetate (1:3 v/v) at −20°C. To serve as a control, the initial ‘input’ cell pellet was re-suspended in 100 µl of 50 µM NaOH and incubated at 95°C for 30 min, followed by pH neutralization by the addition of 10 µl of 1M Tris, pH8. Standard PCR assays targeting a 795 bp chromosomal DNA amplicon were performed to assess bacterial DNA associated with immunopurified CagY complexes. Quantitation of DNA amplification from ‘input’ and ‘IP’ samples was conducted by densitometry analysis in Fiji^79^ and efficiency of ‘IP’ sample amplification was calculated as a percent of the corresponding ‘input’ sample amplification for each biological replicate experiment.

### Bacterial adenylate cyclase two-hybrid (BACTH) assay

To screen direct protein-protein interactions, we used a bacterial two-hybrid system in which the bait protein of interest was fused to the either the T18 N-terminus (pUT18C vector) or the T25 N-terminus (pKNT25 vector) as previously described^62, 80–82^. Plasmids encoding various prey proteins were cloned into both the pUT18C and pKNT25 vectors. Recombinant plasmids were verified by whole plasmid DNA sequencing. Both bait and prey proteins were cloned into *E. coli* BTH101 (Δ*cya*) and transformants were selected on LB plates supplemented with 100 μg/mL ampicillin and 50 μg/mL kanamycin at 30°C. Empty T18 and T25 vectors were co-transformed to serve as a negative control while T18-Zip/T25-Zip^62^ and T18-DprA/T25-DprA^64^ were co-transformed to serve as positive controls. Single colonies were selected, grown overnight at 30°C in LB supplemented with dual antibiotics (100 μg/mL ampicillin and 50 μg/mL kanamycin) and 0.5 mM IPTG, and spotted onto LB plates supplemented with dual antibiotics, 0.5mM IPTG, and 40 mg/mL X-gal. Plates were monitored for changes in colony color (blue indicating positive protein-protein interaction, white indicating negative protein-protein interactions). Growth in M63-0.4% maltose was used to confirm protein-protein interactions. For these studies, 2.5 μL of LB overnight cultures were inoculated into fresh M63-maltose minimal media and were grown under aerobic conditions at 30°C for 72 hours. Bacterial growth was determined by measuring the OD_600_ (Biotek Synergy H1) and protein-protein interactions were compared to the positive and negative control strains. Culture growth was expressed as the fold change in OD_600_ over the corresponding negative control culture (BTH101 co-transformed with empty vectors), and data represent a minimum of at least three independent biological replicate experiments for each pairwise co-transformation.

### Recombinant protein purification

To generate maltose binding protein (MBP)-tagged recombinant fusion proteins, genes of interest were amplified from *H. pylori* 26695 and cloned into pMAL-c6T (New England Biolabs) for protein expression and purification via affinity chromatography as previously described^83^. Briefly, MBP fusions were expressed in Lemo21 (DE3) *E. coli* by induction with 0.5 mM IPTG for 3 hours at 37°C. Bacteria were collected via centrifugation and lysed by sonication in lysis buffer (20 mM Tris pH 7.5, 500 mM NaCl, 1 mM EDTA, 5 mM BME, 1X protease inhibitor cocktail) followed by centrifugation at 16000 x g for 20 min at 4°C. Soluble MBP fusions were passed through a column containing equilibrated amylose resin 3 times at a low flow rate. Columns were washed to remove unbound protein fractions using one column volume of lysis buffer (20 mM Tris, pH 7.5, 500 mM NaCl, 1 mM EDTA, 5 mM BME), one column volume of high salt lysis buffer (20 mM Tris, pH 7.5, 1 M NaCl, 1 mM EDTA, 5 mM BME), followed by a final wash with one column volume lysis buffer. MBP fusions were eluted in 10 mM maltose, 20 mM Tris, pH 7.5, 100 mM NaCl, 1 mM EDTA, 5 mM BME. Protein size and purity was determined by SDS-PAGE analysis and Coomassie staining, and protein concentration was determined by Bradford assay (Bio-Rad). Purified MBP fusions were stored in elution buffer at −80°C.

### DNA binding electromobility shift assays (EMSA)

5′ Cy5-labeled chemically synthesized oligonucleotides containing the *tfs3 oriT*-like recognition sites^83^ were incubated with varying concentrations of MBP fusion proteins ranging from 0 µM to 8 µM. DNA binding reactions were assembled in 15 µl total volume containing 0.01 μM DNA substrate in reaction buffer (50 mM Tris-HCl pH 8, 50 mM KCl, 0.5 mM MgCl_2_, 1 mM DTT, 0.1 µg/µl BSA) for 30 min at 4°C. Free DNA and nucleoprotein complexes were resolved using native PAGE (10%) run in 1X TBE at 70V for 2-3h. For competition assays, unlabeled competitor oligonucleotides (0.01 μM to 0.4 μM) were added to the reaction mixture containing Cy5-labeled oligonucleotides (0.01 μM) and MBP-CagX (0.75 μM). After incubation on ice for 30 min, reaction mixtures were resolved using native PAGE (10%) run in 1X TBE at 70V for 2-3h at 4°C. Quantification of free DNA was performed using ImageLab software (Bio-Rad).

### Protein structure modeling

CagC predicted structure was determined using AlphaFold2 colab^54, 55^. Protein sequence alignments were performed using MegAlign Pro (DNAstar). Superimposed images of protein structures were generated using ChimeraX Matchmaker^84^. Structural analyses of the *H. pylori cag* T4SS PRC were conducted on PDB 6X6J. The electrostatic surfaces of CagX, TraK, and VirB9 were generated as coulombic potential maps in ChimeraX.

### Immunoblotting

To detect bacterial protein expression, normalized samples were lysed in reducing 2X SDS buffer (Bio-Rad), resolved by SDS-PAGE (10%), transferred to a nitrocellulose membrane, and immunoblotted using rabbit polyclonal antisera raised against CagC peptides^69^ or recombinant CagX (CagC and CagX antisera were a gift from Dr. Tim Cover) as previously described^48^. To confirm equal sample loading, membranes were immunoblotted using *H. pylori* outer membrane protein monoclonal antibody (α-OMP, Santa Cruz) or anti-*H. pylori* polyclonal antisera raised against *H. pylori* whole cell lysate (a gift from Dr. Tim Cover). Secondary detection by horse radish peroxidase-conjugated anti-rabbit IgG or anti-mouse IgG was achieved using SuperSignal West Pico chemiluminescent substrate (Thermo).

### IL-8 quantitation

Levels of IL-8 secreted by AGS cells co-cultured with *H. pylori* or corresponding isogenic derivatives were determined using the human CXCL8 ELISA (R&D Systems). Briefly, AGS cells were challenged with *H. pylori* or the indicated isogenic mutant strain for 4.5 h at an MOI of 100. Supernatants were collected and analyzed as previously described^48, 49^. Biological replicate experiments were performed a minimum of three times.

## ACKNOWLEDGMENTS

Work in the Shaffer lab is funded by the NIH (P20 GM130456 to CLS) and academic development funds provided by the University of Kentucky (to CLS).

## FIGURE LEGENDS

**Supplemental Figure 1.**
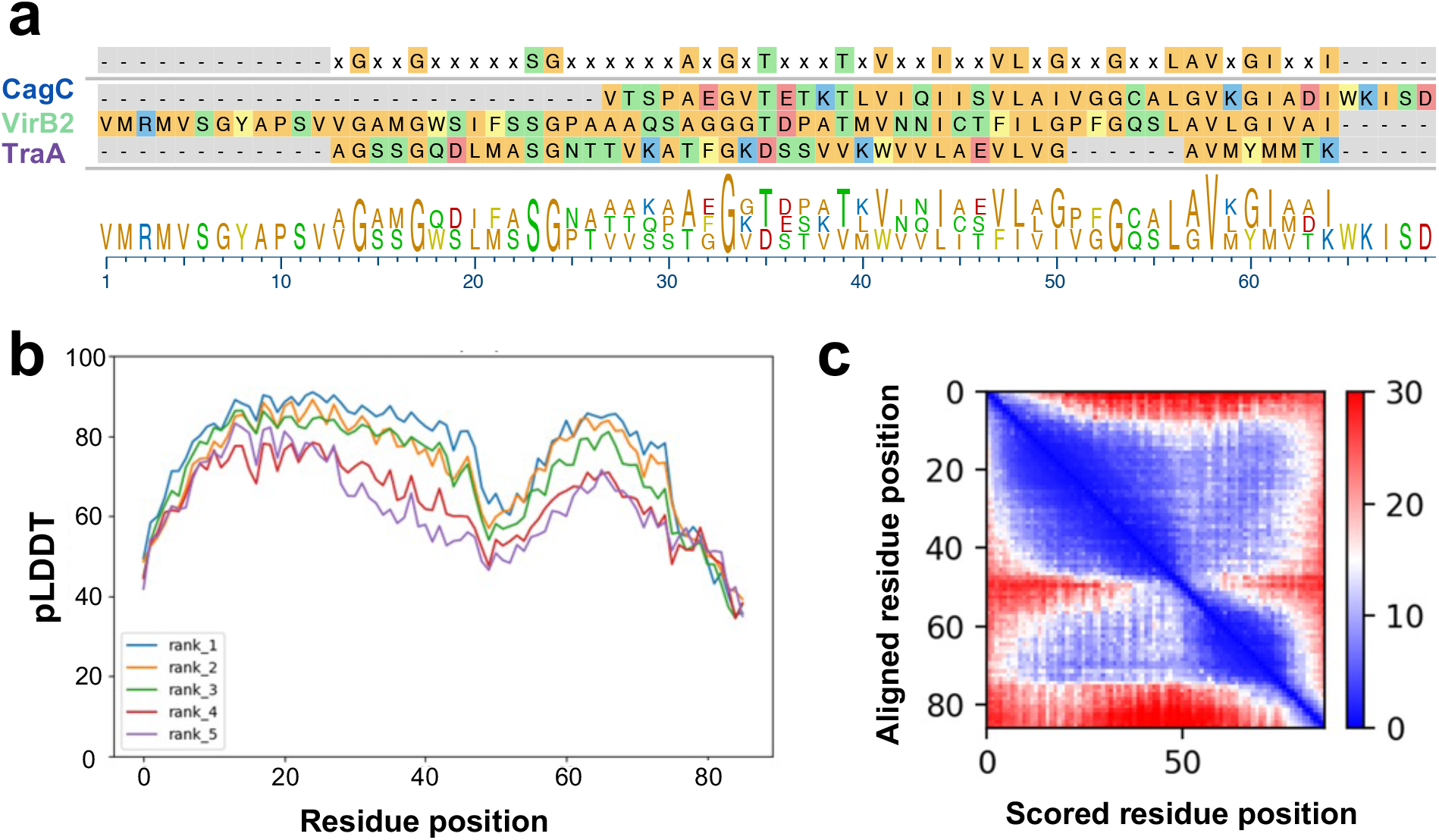
CagC structural modeling. (**a**) Protein sequence alignment of prototypical T4SS pilins VirB2 (*A. tumefaciens vir* T4SS), TraA (*E. coli* F pilus pED208), and CagC. Pilin sequence alignment was generated using MegAlign Pro. **(b**) Modeling scores determined by AlphaFold2 for the highest-ranked CagC models (pLDDT, predicted local distance difference test, representing local model accuracy). (**c**) Heatmap representing the inter-chain accuracy of the highest-rank CagC model. Heatmap pixels indicate alignment error (in Angstroms) for a pair of two residues, indicating the positional uncertainty at residue x if the predicted and actual structures are aligned on residue y^56^.

**Supplemental Figure 2.**
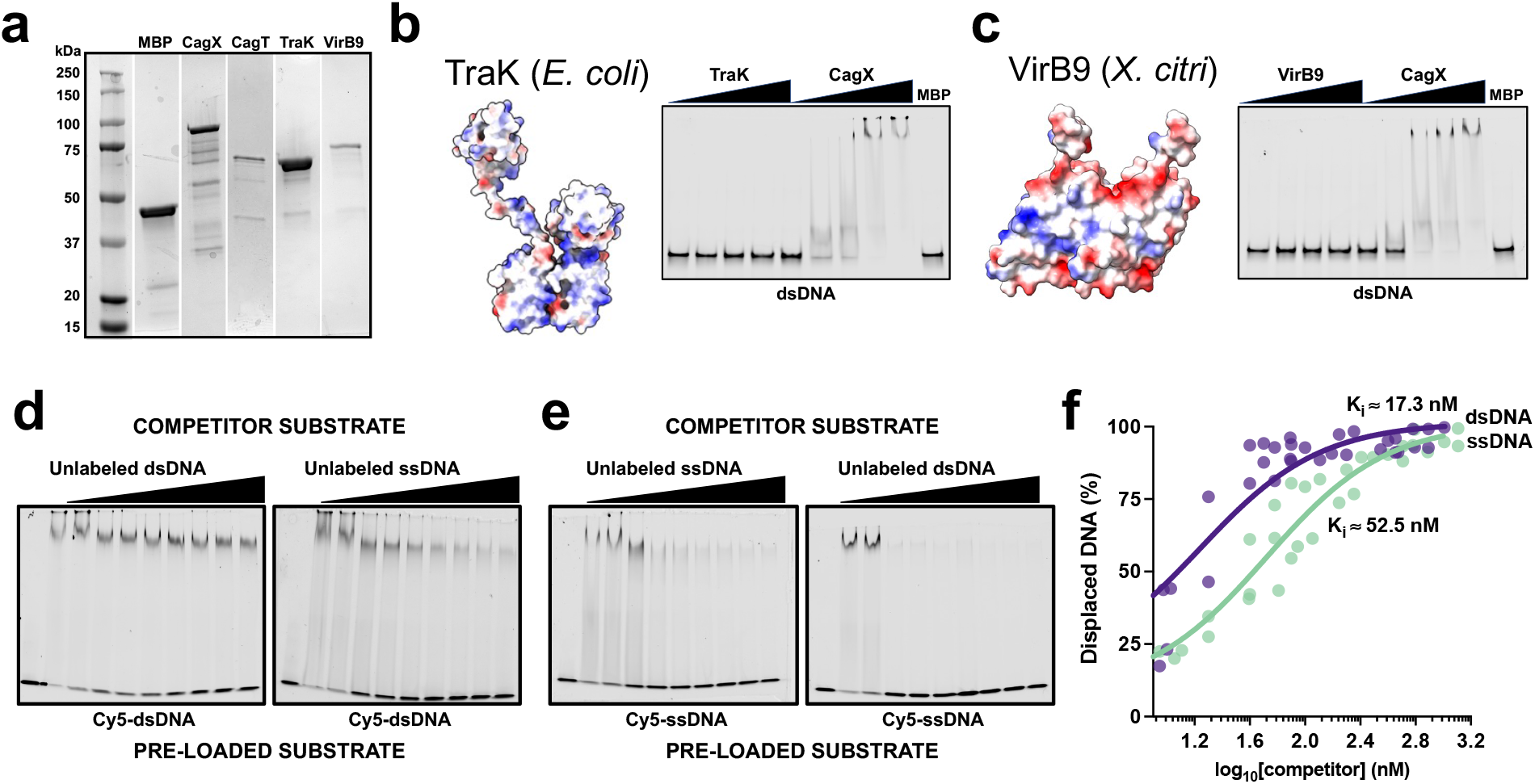
CagX preferentially binds target dsDNA. (**a**) Coomassie analysis of recombinant maltose binding protein (MBP) fusions purified from *E. coli*. (**b**) 58-mer dsDNA EMSA analysis of TraK derived from F plasmid R1 (structural model depicts TraK from F plasmid pED208, PDB: 7SPB) and (**c**) VirB9 derived from *A. tumefaciens vir* T4SS (structural model depicts VirB9 from *X. citri vir* T4SS, PDB: 6GYB). MBP-CagX, but not MBP-TraK or MBP-VirB9, binds dsDNA. (**d, e**) Competitive DNA binding assays demonstrating CagX preferential dsDNA binding. Recombinant CagX was pre-loaded with Cy5-labeled dsDNA (**d**) or ssDNA (**e**) and the indicated cold competitor was added at increasing concentrations. (**f**) Quantitation of displaced DNA in Cy5-ssDNA pre-loaded reactions. Data points represent the % displaced DNA obtained in independent biological replicate experiments. Solid line represents the non-linear regression using the one site fit K_i_ model in GraphPad Prism 9.5 to estimate the indicated inhibitory constants (K_i_) for dsDNA and ssDNA.

**Supplemental Figure 3.**
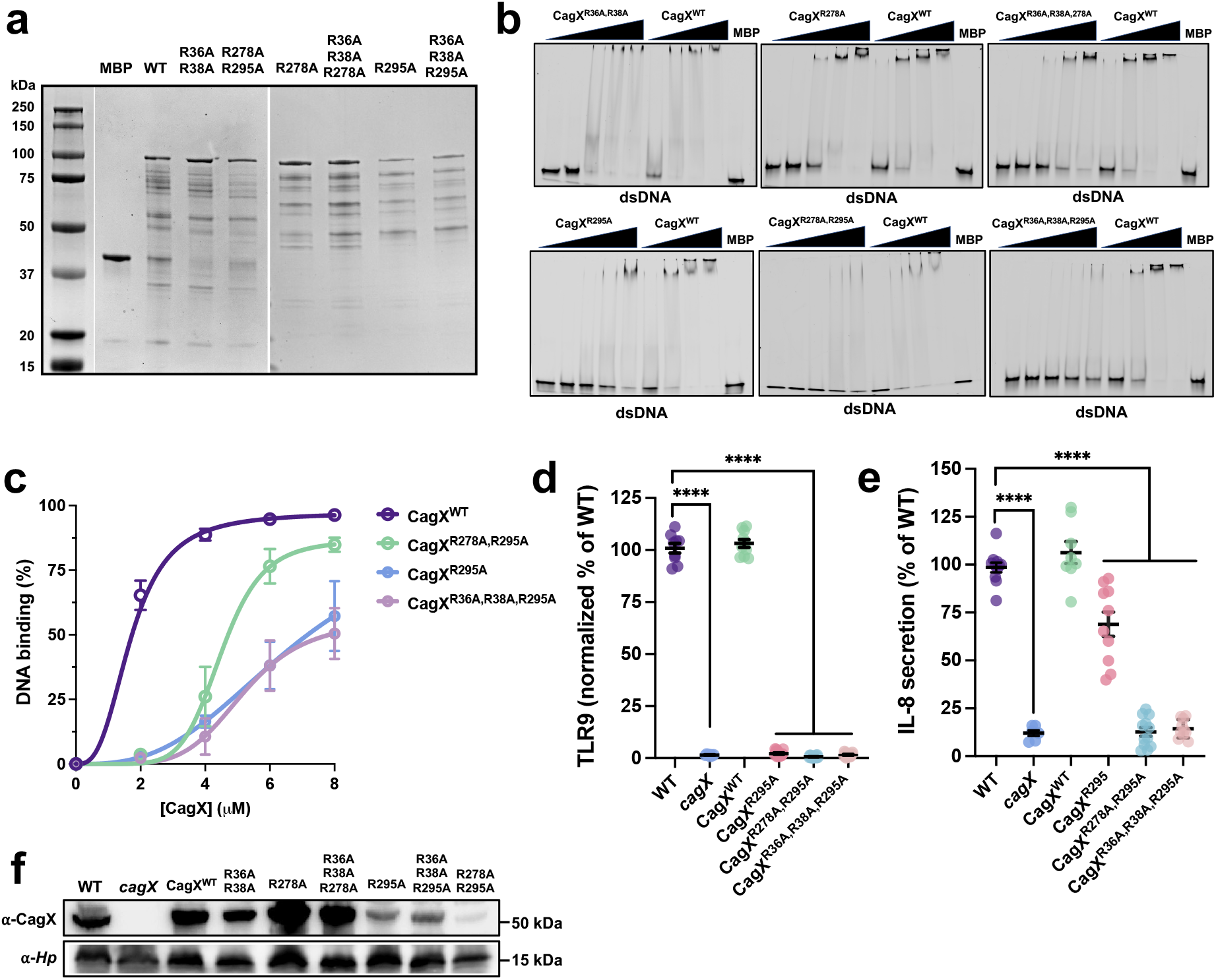
Intersubunit π-stacking is required for CagX dsDNA binding. (**a**) Coomassie analysis of recombinant maltose binding protein (MBP) CagX variant fusions purified from *E. coli*. (**b**) EMSA analyses of recombinant CagX variant binding to dsDNA. (**c**) Quantitation of MBP-CagX binding to 58-mer DNA substrates. The experimentally approximated K_D_ for CagX- DNA binding was determined for CagX^WT^ (1.7 µM), CagX^R278A,R295A^ (4.5 µM), CagX^R95A^ (6.9 µM), and CagX^R36A,R38A,R295A^ (5.2 µM). (**d**) TLR9 stimulation and (**e**) IL-8 secretion induced by *H. pylori* strains. (**f**) Levels of CagX produced by *H. pylori cagX* harboring the indicated CagX variant. In **c**, dissociation constants (K_D_) were estimated using the specific binding with hill slope nonlinear regression model in GraphPad Prism 9.5. In **d** and **e**, significance was determined by one-way ANOVA with Dunnett’s post-hoc correction for multiple comparisons to controls; ****, *p*<0.0001.

